# Analyzing Differences in Processing Nouns and Verbs in the Human Brain using Combined EEG and MEG Measurements

**DOI:** 10.1101/2024.12.04.626813

**Authors:** Nikola Kölbl, Nadia Müller-Voggel, Stefan Rampp, Martin Kaltenhäuser, Konstantin Tzirdis, Patrick Krauss, Achim Schilling

## Abstract

Language and consequently the ability to transmit and spread complex information is unique to the human species. The disruptive event of the introduction of large language models has shown that the ability to process language alone leads to incredible abilities and, to some extent, to intelligence. However, how language is processed in the human brain remains elusive. Many insights originate from fMRI studies, as the high spatial resolution of fMRI devices provides valid information about where things happen. Nevertheless, the limited temporal resolution prevents us from gaining a deep understanding on the underlying mechanisms. In this study, we performed combined EEG and MEG measurements in 29 healthy right-handed human subjects during the presentation of continuous speech. We compared the evoked potentials (ERPs and ERFs) for different word types in source space and sensor space across the whole brain. We found characteristic spatio-temporal patterns for different word types (nouns, verbs) especially at latencies of 300ms to 1 s. This is further emphasized by the fact that we observe these effects in two pre-defined sub-samples of the data set (exploration and validation sample). We expect this study to be the starting point for further evaluations of semantic and syntactic processing in the brain.

## Introduction

Despite the fact that certain animal species such as toothed whales have complex communication systems (Bergler et al., 2019; Janik, 2014), humans are the only species with complex language which could be used to express an infinite amount of ideas and information (Berwick, Friederici, Chomsky, & Bolhuis, 2013). The fact that language and general intelligence are tightly linked is emphasized by the fact that already Alan Turing developed the “Imitation Game”-a language-based task to test intelligence of computers (Turing, 1950; Moor, 1976; Piccinini, 2000). Despite the fact that language might be one key to general intelligence, as proposed by research on Large Language Models (LLMs) (Bubeck et al., 2023), language processing in the human brain is not well understood. Thus, through high-resolution fMRI measurements we know the main regions of language processing in the human brain such as Broca’s area, Wernicke’s area or the arcuate fasciculus (Goucha & Friederici, 2015; Friederici, Meyer, & Von Cramon, 2000; Friederici, 2015), however the exact mechanisms behind language processing in the brain stay elusive. To further investigate the mechanisms, methods with higher temporal resolution such as electroencephalography (EEG), invasive EEG (Metzger et al., 2020) and magnetoencephalography (MEG) (Tavabi, Obleser, Dobel, & Pantev, 2007) are needed. Recently, in neurolinguistic research, the trend has shifted from presenting repetitively the same single artificial stimuli, such as single sentences under very controlled conditions (see e.g. (Praamstra & Stegeman, 1993)), to presenting natural continuous speech such as audio books (Schilling et al., 2021; Koelbl, Schilling, & Krauss, 2023; Garibyan, Schilling, Boehm, Zankl, & Krauss, 2022; Schüller, Schilling, Krauss, Rampp, & Reichenbach, 2023; Schüller, Schilling, Krauss, & Reichenbach, 2024). In the present study, we played the beginning of two story lines (corresponds to 6.5 chapters) of the German science fiction audio book “Vakuum” from Phillip P. Peterson to 29 healthy right-handed participants and performed combined MEG and EEG measurements. Furthermore, we extracted event-related-fields (ERFs) and event-related-potentials (ERPs) from the continuous data stream. We found distinct spatio-temporal patterns for different wordtypes (nouns, verbs). These findings were consistent across two independent sub-samples of the dataset, defined before evaluation (exploration and validation set), analogous to training and test datasets in Machine Learning (Schilling et al., 2021).

### Data

In this study, 29 participants (15 ♀, average age: 22.8 ± 3 years) listened to the German audio book “Vakuum” by Philip P. Peterson (Argon Verlag) split in eight sections (approx. 50 minutes). All were healthy, righthanded native German speakers. The brain activity was recorded simultaneously using 248 MEG and 64 EEG channels. The study was approved by the Ethics Committee of the University Hospital Erlangen (Approval No: 22-361-2, PK). The data were pre-processed identically for EEG and MEG. We identified and interpolated sensors with poor signal-to-noise ratios, applied a 1-20 Hz bandpass filter, sampled the data down to 200Hz and did a further ICA-based filtering step for artefact removal (according to (Koelbl et al., 2023)). Finally, the data was segmented into intervals from -0.5 s before to 1 s after each word onset. The data was split into exploration (subjects 1-19) and validation (subjects 20-29) sets prior to the evaluation, in order to check for consistency. This data splitting is analogous to AI approaches, where data is split into a training and test data set.

Grand Average ERPs/ERFs were calculated for both data sets for three latency-intervals: 0.0-0.3 s, 0.3-0.6 s and 0.6-0.9 s (Figure 1). To assign the respective signals to brain regions, we performed source localization using a cortical volume source space and the boundary element model of the template brain *fsaverage* from *FreeSurfer* (https://surfer.nmr.mgh.harvard.edu/). We first computed the inverse operator using the noise covariance matrices from the pre-stimulus intervals and the previously computed forward solution. Then, by applying this operator to the average evoked data with method=“sLORETA”, we generated a volume vector source estimate (SE). This process is repeated for both word types and sets and the grand average time-averaged SEs of the validation set in the middle and last interval are visualized in Figure 2.

**Figure 1:**
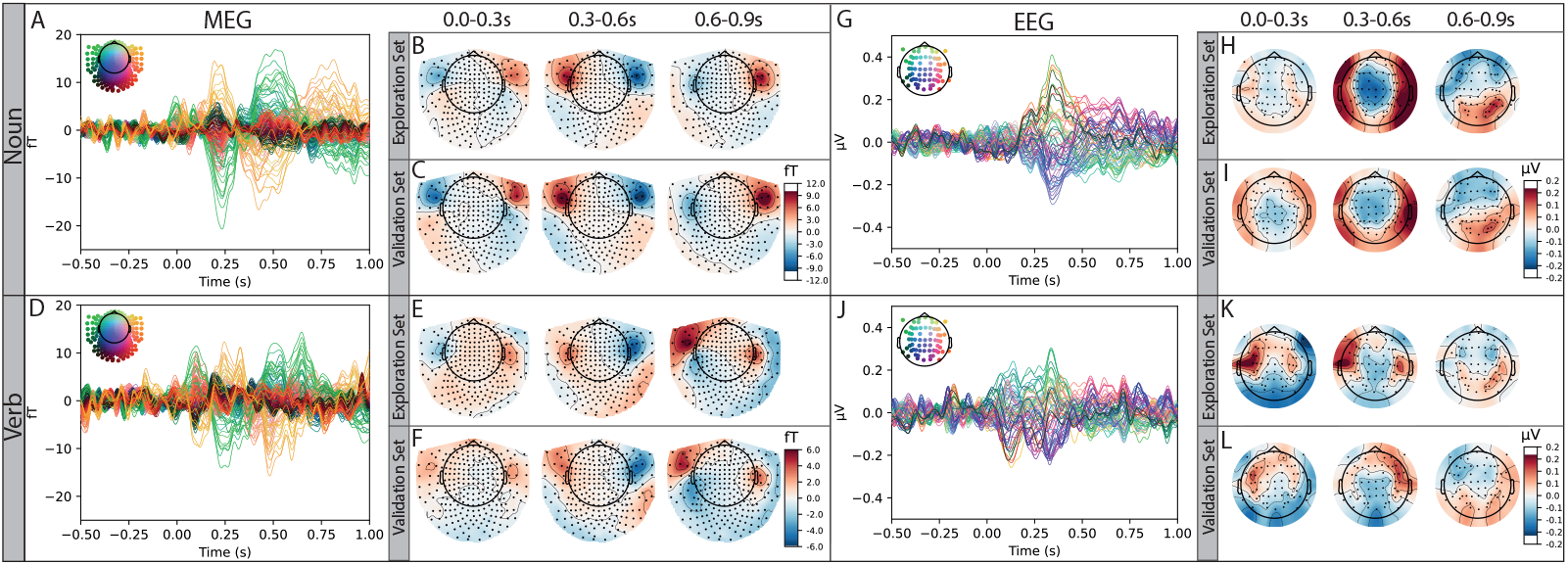
ERFs and ERPs in source space. The figure shows the ERFs as a response to the presentation of nouns (A) and verbs (D). The temporal activity is drawn as topomaps (nouns: B, C; verbs: E, F). Before evaluation we have split the data into a exploration data set (B, E) and a validation set (C, F) in order to test if the effects are reproducible observed. G-J: Is the same analysis as A-F but for the simultaneously recorded EEG data.

**Figure 2:**
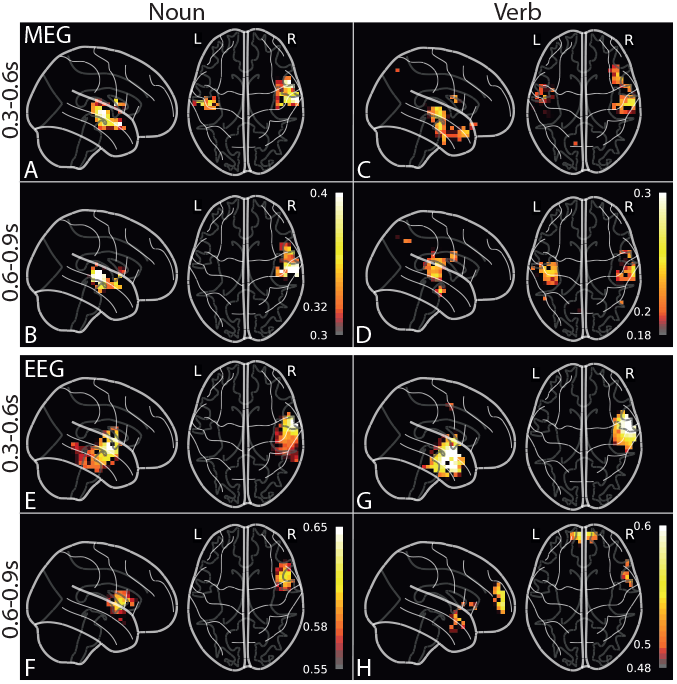
Evoked activity in sensor space. We averaged the activity across two different latency ranges (300-600 ms and 600-900 ms) and found different activity patterns for verb and nouns.

## Results

We found clear differences in the ERFs (Fig. 1 A-F) and the ERPs (Fig. 1 G-L) evoked by the presentation of nouns compared to verbs. The spatio-temporal activity is highly variable for the different word types, but consistent over the exploration and validation data set (subsamples of the data defined before the evaluation). The distinct activity in the right hemisphere (see Fig. 2) suggests that the differences between word types are not exclusively caused by different syntactic processing, but may be due to differences in the phonetic characteristics of the words or contextual information.

## Discussion

By analyzing two independent sub-samples (exploration, validation) of our MEG/EEG data set, we showed that continuous speech stimuli can be used to identify different cortical activity patterns related to the processing of different word types. To disentangle the effects caused by semantic, syntactic, and phonetic processing in the brain, it is important to further investigate the data with innovative (e.g. AI-based) evaluation techniques (see e.g. (Schilling et al., 2022; Metzner, Schilling, & Krauss, 2023; Metzner et al., 2022; Krauss et al., 2018, 2021)) and to combine the measurements with a solid theoretical background and thus, to follow the cognitive-computational-neuroscience philosophy (CCN, (Kriegeskorte & Douglas, 2018)) proposing to use biologically plausible AI algorithms as model for the brain (see also (Schilling, Sedley, et al., 2023; Schilling, Schaette, et al., 2023; Gerum, Erpenbeck, Krauss, & Schilling, 2020; Gerum & Schilling, 2021)).

## Acknowledgments

This work was funded by the Deutsche Forschungsgemeinschaft (DFG, German Research Foundation): grants KR 5148/2-1 (project number 436456810), KR 5148/3-1 (project number 510395418) and GRK 2839 (project number 468527017) to PK, grant TZ 100/2-1 (project number 510395418) to KT, and grant SCHI 1482/3-1 (project number 451810794) to AS.

## References

Bergler, C., Schröter, H., Cheng, R. X., Barth, V., Weber, M., Nöth, E., … Maier, A. (2019). Orcaspot: An automatic killer whale sound detection toolkit using deep learning. Scientific reports, 9(1), 10997.

Berwick, R. C., Friederici, A. D., Chomsky, N., & Bolhuis, J. J. (2013). Evolution, brain, and the nature of language. Trends in cognitive sciences, 17 (2), 89–98.

Bubeck, S., Chandrasekaran, V., Eldan, R., Gehrke, J., Horvitz, E., Kamar, E., … others (2023). Sparks of artificial general intelligence: Early experiments with gpt-4. arXiv preprint 2303.12712.

Friederici, A. D. (2015). White-matter pathways for speech and language processing. Handbook of clinical neurology, 129, 177–186.

Friederici, A. D., Meyer, M., & Von Cramon, D. Y. (2000). Auditory language comprehension: an event-related fmri study on the processing of syntactic and lexical information. Brain and language, 74(2), 289–300.

Garibyan, A., Schilling, A., Boehm, C., Zankl, A., & Krauss, P. (2022). Neural correlates of linguistic collocations during continuous speech perception. Frontiers in Psychology, 13, 1076339.

Gerum, R. C., Erpenbeck, A., Krauss, P., & Schilling, A. (2020). Sparsity through evolutionary pruning prevents neuronal networks from overfitting. Neural Networks, 128, 305–312.

Gerum, R. C., & Schilling, A. (2021). Integration of leaky-integrate-and-fire neurons in standard machine learning architectures to generate hybrid networks: A surrogate gradient approach. Neural Computation, 33(10), 2827–2852.

Goucha, T., & Friederici, A. D. (2015). The language skeleton after dissecting meaning: A functional segregation within broca’s area. NeuroImage, 114, 294–302.

Janik, V. M. (2014). Cetacean vocal learning and communication. Current opinion in neurobiology, 28, 60–65.

Koelbl, N., Schilling, A., & Krauss, P. (2023). Adaptive ica for speech eeg artifact removal. In 2023 5th international conference on bio-engineering for smart technologies (biosmart) (pp. 1–4).

Krauss, P., Metzner, C., Joshi, N., Schulze, H., Traxdorf, M., Maier, A., & Schilling, A. (2021). Analysis and visualization of sleep stages based on deep neural networks. Neurobiology of sleep and circadian rhythms, 10, 100064.

Krauss, P., Metzner, C., Schilling, A., Tziridis, K., Traxdorf, M., Wollbrink, A., … Schulze, H. (2018). A statistical method for analyzing and comparing spatiotemporal cortical activation patterns. Scientific reports, 8(1), 5433.

Kriegeskorte, N., & Douglas, P. K. (2018). Cognitive computational neuroscience. Nature neuroscience, 21(9), 1148–1160.

Metzger, B. A., Magnotti, J. F., Wang, Z., Nesbitt, E., Karas, P. J., Yoshor, D., & Beauchamp, M. S. (2020). Responses to visual speech in human posterior superior temporal gyrus examined with ieeg deconvolution. Journal of Neuroscience, 40(36), 6938–6948.

Metzner, C., Schilling, A., & Krauss, P. (2023). Beyond labels: Advancing cluster analysis with the entropy of distance distribution (edd). arXiv preprint 2311.16621.

Metzner, C., Schilling, A., Traxdorf, M., Tziridis, K., Maier, A., Schulze, H., & Krauss, P. (2022). Classification at the accuracy limit: facing the problem of data ambiguity. Scientific Reports, 12(1), 22121.

Moor, J. H. (1976). An analysis of the turing test. Philosophical Studies: An International Journal for Philosophy in the Analytic Tradition, 30(4), 249–257.

Piccinini, G. (2000). Turing’s rules for the imitation game. Minds and Machines, 10(4), 573–582.

Praamstra, P., & Stegeman, D. F. (1993). Phonological effects on the auditory n400 event-related brain potential. Cognitive Brain Research, 1(2), 73–86.

Schilling, A., Gerum, R., Boehm, C., Rasheed, J., Metzner, C., Maier, A., … Krauss, P. (2022). Deep learning based decoding of local field potential events. bioRxiv, 2022–10.

Schilling, A., Schaette, R., Sedley, W., Gerum, R. C., Maier, A., & Krauss, P. (2023). Auditory perception and phantom perception in brains, minds and machines. Frontiers in Neuroscience, 17, 1293552.

Schilling, A., Sedley, W., Gerum, R., Metzner, C., Tziridis, K., Maier, A., … Krauss, P. (2023). Predictive coding and stochastic resonance as fundamental principles of auditory phantom perception. Brain, 146(12), 4809–4825.

Schilling, A., Tomasello, R., Henningsen-Schomers, M. R., Zankl, A., Surendra, K., Haller, M., … Krauss, P. (2021). Analysis of continuous neuronal activity evoked by natural speech with computational corpus linguistics methods. Language, Cognition and Neuroscience, 36(2), 167–186.

Schüller, A., Schilling, A., Krauss, P., Rampp, S., & Reichenbach, T. (2023). Attentional modulation of the cortical contribution to the frequency-following response evoked by continuous speech. Journal of Neuroscience, 43(44), 7429–7440.

Schüller, A., Schilling, A., Krauss, P., & Reichenbach, T. (2024). The early subcortical response at the fundamental frequency of speech is temporally separated from later cortical contributions. Journal of Cognitive Neuroscience, 36(3), 475–491.

Tavabi, K., Obleser, J., Dobel, C., & Pantev, C. (2007). Auditory evoked fields differentially encode speech features: an meg investigation of the p50m and n100m time courses during syllable processing. European journal of neuroscience, 25(10), 3155–3162.

Turing, A. M. (1950). Computing machinery and intelligence (1950).

